# A recombinant selective drug-resistant *M. bovis* BCG enhances the bactericidal activity of a second-line tuberculosis regimen

**DOI:** 10.1101/2021.03.04.434024

**Authors:** Gift Chiwala, Zhiyong Liu, Julius N. Mugweru, Bangxing Wang, Shahzad Akbar Khan, Petuel Ndip Ndip Bate, Buhari Yusuf, H.M. Adnan Hameed, Cuiting Fang, Yaoju Tan, Ping Guan, Jinxing Hu, Shouyong Tan, Jianxiong Liu, Nanshan Zhong, Tianyu Zhang

## Abstract

Drug-resistant tuberculosis (DR-TB) results from infection by *Mycobacterium tuberculosis* strains resistant to at least rifampin or isoniazid. To improve the treatment outcome in DR-TB, therapeutic vaccines are considered an ideal choice as they can enhance pathogen clearance and minimize disease sequelae. To date, there is no therapeutic vaccine reported to be effective when combined with a chemotherapy regimen against DR-TB. The only available TB vaccine, the *M. bovis* BCG (BCG) is susceptible to several anti-TB drugs hence not a perfect option for therapeutic vaccination. Herein, we developed a recombinant BCG (RdrBCG) overexpressing Ag85B and Rv2628 with resistance to selected anti-TB drugs. When administered three times adjunct to a second-line anti-TB regimen in a classical murine model of DR-TB, the RdrBCG lowered lung *M. tuberculosis* colony-forming units by 1 log_10_. Furthermore, vaccination with the RdrBCG adjunct to TB chemotherapy minimized lung tissue pathology in mice. Most importantly, the RdrBCG maintained the exogenously inserted genes and showed almost the same virulence as its parent BCG Tice strain in severe combined immune-deficient mice. All these suggested that the RdrBCG was stable, safe and effective. Hence, the “recombinant” plus “drug-resistant” BCG strategy could be a useful concept for developing therapeutic vaccines against DR-TB.

## Introduction

Tuberculosis (TB) is a chronic infectious disease caused by *Mycobacterium tuberculosis* (*M. tuberculosis*) and a serious threat to public health for it is the leading cause of morbidity and mortality around the globe (1). In 2019, 10 million people developed TB disease and about 1.4 million people died from infection with TB globally. Drug-resistant (DR), especially the multiple DR (MDR)-TB or even extensive DR (XDR)-TB, caused by drug-resistant *M. tuberculosis* strains continue to be a public health crisis responsible for approximately half a million cases in 2019 (1, 2). MDR-TB is resistant to at least isoniazid and rifampin and unfortunately, only limited drug options are available with low cure rates (3). Furthermore, its treatment duration is very long (9 to 24 months for currently available regimens) which at times results in treatment interruptions or default thereby causing more drug resistance. The situation is even worse for XDR-TB which is MDR-TB with resistance to a fluoroquinolone and a second-line injectable agent as it requires multiple drug combinations administered over a long time (4).

Prevention of TB depends on a rapid and timely diagnosis as well as treatment of the patient(s) until they are rendered non-infectious and disease free. Additional strategies include the treatment of persons with latent TB infection who are at risk of developing active disease and vaccination with *Mycobacterium bovis* bacille Calmette Guerin (BCG) (1, 5). Recently, there has been tremendous interest in the use of therapeutic vaccine(s) as well as host-directed therapies to stimulate the host’s immune response whilst on drug therapy to improve the treatment outcome (6, 7). To date, several newly developed anti-TB vaccines like the recombinant BCG vaccine VPM1002 or *Mycobacterium vaccae,* have entered clinical trials with some still in the preclinical phases (8–10). Although BCG is effective in preventing severe forms of TB in children, its use as a therapeutic vaccine would easily fail as it is susceptible to the administered anti-TB drugs (11–13). There is, therefore, a need to find a vaccine that will not only stimulate the required and appropriate immune response but also enhance the *M. tuberculosis* killing efficiency while resisting the administered anti-TB drugs.

Antigen (Ag) 85B is a member of the Ag85 complex which comprises of Ag85A, Ag85B and Ag85C (14). These antigens exhibit enzymatic activity as mycolyl transferases and catalyze transesterification reactions leading to the synthesis of trehalose monomycolate, trehalose dimycolate (TDM), and mycolated arabinogalactan (15). TDM and trehalose monomycolate form part of the mycobacteria’s cell wall and play important roles in recruiting cells for granuloma formation and modulates the expression of immune mediators. Additionally, TDM enhances *M. tuberculosis* replication and viability thereby contributing to the persistence of *M. tuberculosis* infection (16, 17). The mycolated arabinogalactan is covalently linked to the peptidoglycan layer and is also essential for cell viability (18). Reportedly, a recombinant BCG co-expressing Ag85B, ESAT-6 and Rv2608 induced a strong CD4^+^ and CD8^+^ T-lymphocytes proliferation in vaccinated mice than the wild-type controls. Furthermore, this recombinant BCG elicited more Th1 and humoral immune response but suppressed IL-10 expression (19). Th1 response is essential for activating macrophage associated killing of intracellular pathogens including *M. tuberculosis* by enhancing phagolysosome fusion (20, 21).

The use of latency antigens which are expressed under stressful conditions like hypoxia is also being explored in developing effective anti-TB vaccines (22). Although the functions of most latency antigens or DosR regulon proteins are not known, evidence shows that they aid *M. tuberculosis’* survival during times of deprivation like low oxygenation in granulomas (23, 24). Hence they are thought to aid the BCG’s survival within the host macrophages which may improve the potency of BCG against the host defenses (25). Furthermore, the DosR proteins have a strong recognition for T cells as well as IFN-γ which is suggestive of their potential diagnostic or immunogenic role (26). Rv2628 is a putative hypothetical protein induced during hypoxia or stressful conditions and helps the *M. tuberculosis* survive such conditions in the host macrophages and/or granulomas (23, 27). Reportedly, the Rv2628 has a strong T-cell and IFN-γ inducing capacity, which necessitates its exploration for vaccine development against *M. tuberculosis* (28, 29). Most importantly, vaccination of mice with a plasmid expressing Rv2628 induced expression of antigen specific antibody response which is suggestive of its potential role in TB vaccine development (30). Additionally, the Rv2628 enhanced a sustained IgG antibody expression in vaccinated and *M. tuberculosis* challenged mice (31). Despite so many advances in TB vaccine research and development, there’s no vaccine with proven preventive efficacy than the traditional BCG against TB.

The goal of this study was to assess if a recombinant BCG vaccine with resistance to selected anti-TB drugs (RdrBCG) has a therapeutic effect in DR-TB. The approach of using recombinant drug-resistant BCG is novel and allows the BCG’s administration during chemotherapy. Specifically, we selected and verified DR-BCG strains and transformed them with plasmids pEBCG and pIBCG expressing Ag85B and Rv2628. Many studies have reported the immune reactive role that Ag85B and Rv2628 play in stimulating Th1 response which is protective against TB. Hence, we hypothesized that a recombinant BCG vaccine co-expressing Ag85B and Rv2628 with resistance to selected anti-TB drugs would enhance bacterial clearance when co-administered with the corresponding drugs in a murine model 5 of TB. We found that the RdrBCG survived the administered anti-TB drugs and lowered the *M. tuberculosis* bacterial burden as well as minimized pulmonary tissue pathology in vaccinated mice. Furthermore, the RdrBCG showed almost the same virulence as its parent BCG Tice strain in severe combined immune deficient (SCID) mice.

## Results

### DR-BCG harboring mutations in drug resistance-related genes

To obtain DR-BCG strains, we screened for spontaneous DR-mutants using different drugs and routes in parallel. The obtained DR-BCG were verified by drug susceptibility testing (DST) and sequencing of the resistance-related genes in each step. Here we only describe the route by which we successfully obtained the target DR-BCG. Upon screening the wild-type BCG Tice (WtBCG) on streptomycin (S) we obtained three colonies at 10 µg mL^−1^ with a nonsynonymous single nucleotide polymorphism (SNP) encoding Lys43Arg mutation in RpsL (Table 1). When the 3 S-resistant mutants were screened on levofloxacin (L), three colonies resistant to L at 2 µg mL^−1^ were obtained, each harboring a Gly88Asn SNP in the GyrA. Further screening of the 3 S-L-resistant mutants on ethambutol (E) yielded three true mutants resistant to E 5 µg mL^−1^. We obtained several false E-resistant colonies during screening runs whose E minimum inhibitory concentrations (MIC) did not change significantly. This could probably be due to the bacteriostatic but not bactericidal activity of E against BCG. On sequencing, all the three true E-resistant mutants harbored Val49Ile, Glu149Asp nonsynonymous SNPs in Rv3806 and synonymous SNP in EmbB and EmbC (Table 1). We then screened the S-L-E-resistant mutants on prothionamide (P) plates containing 30 µg mL^−1^ and isolated only one mutant with Trp21Arg mutation in EthA. Finally, we plated the S-L-E-P-resistant mutant on 10 µg mL^−1^ amikacin (A) containing plates and ended up with 3 mutants harboring AGG192AG deletion in *rrs* (Table 1).

**Table 1:**
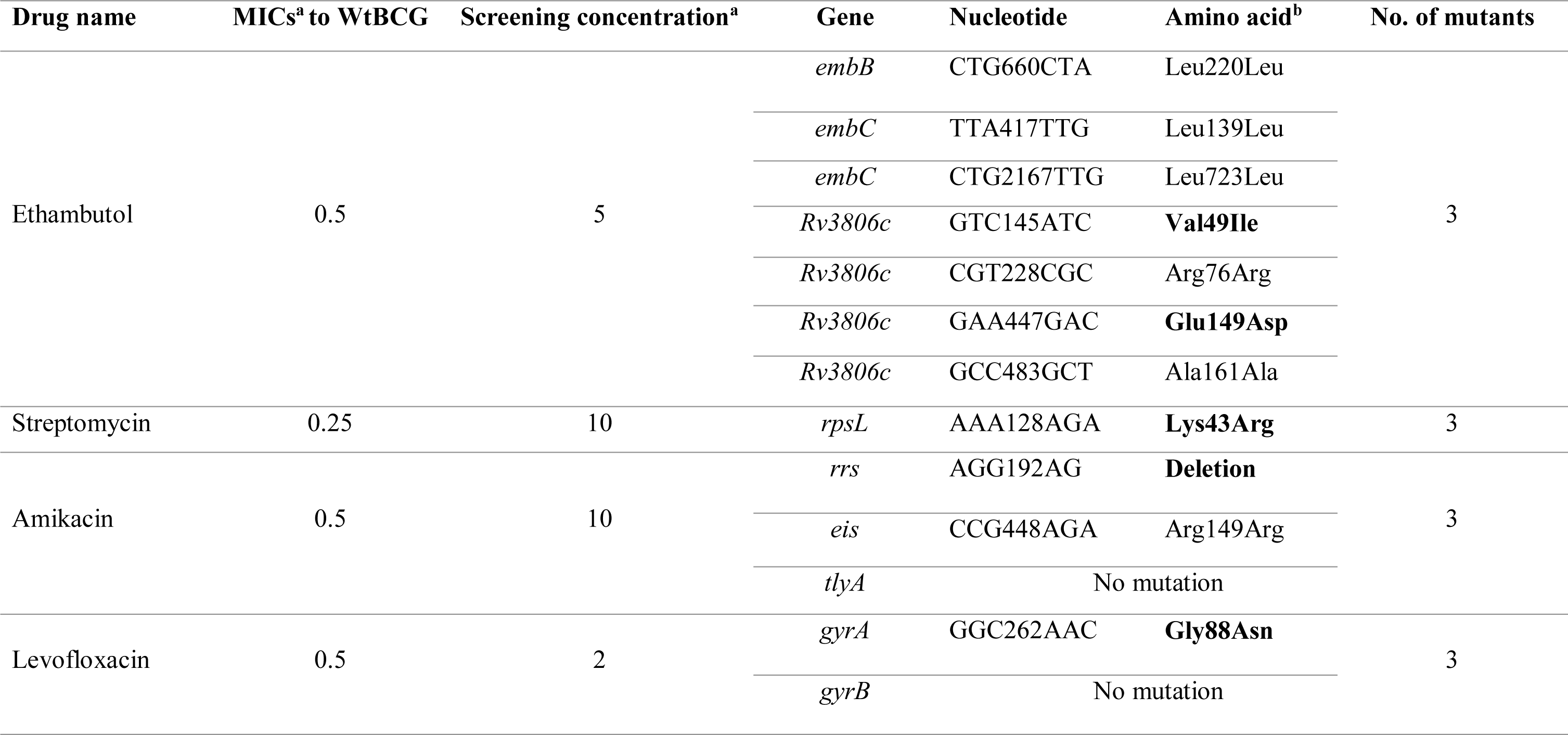
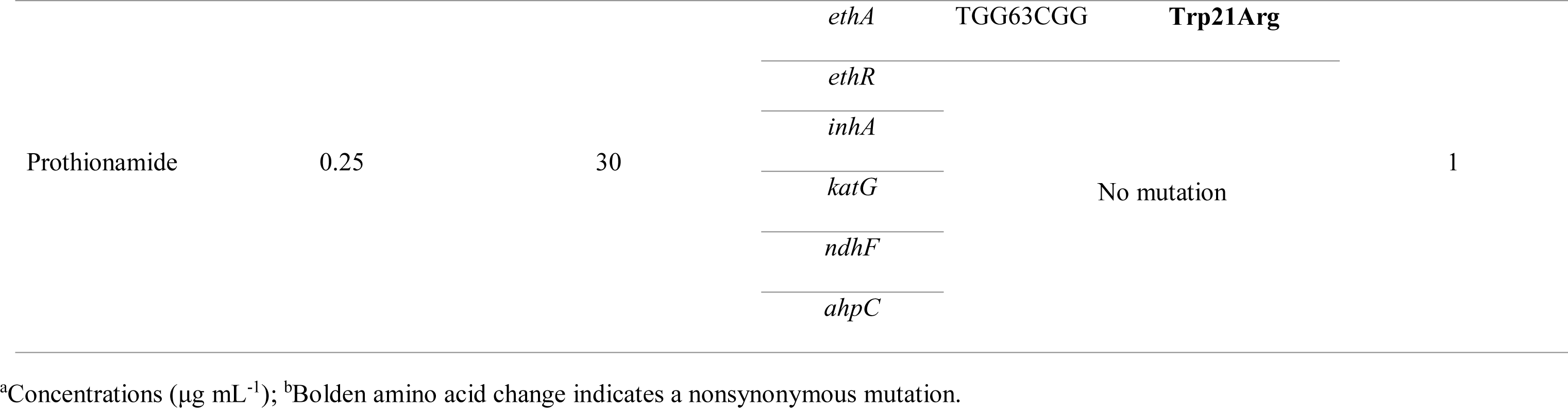
Identification of the corresponding mutations in the selected spontaneous DR-BCG mutants

### Construction of RdrBCGs

To create the RdrBCG strains expressing Ag85B and Rv2628, we first constructed two plasmids, pEBCG and pIBCG (Fig. 1) which were verified by restriction enzyme cut and sequencing. Both plasmids contained the same *hsp60*-*Rv2628-Ag85B* expression cassette. The main difference between the two plasmids is that the pEBCG is a multicopy extra-chromosome plasmid while the pIBCG is a single copy integrative plasmid whose *dif*-*int*-*hyg-dif* cassette containing hygromycin resistance gene (*hyg*) can be removed by the endogenous XerCD recombinases of BCG (32).

**Figure 1:**
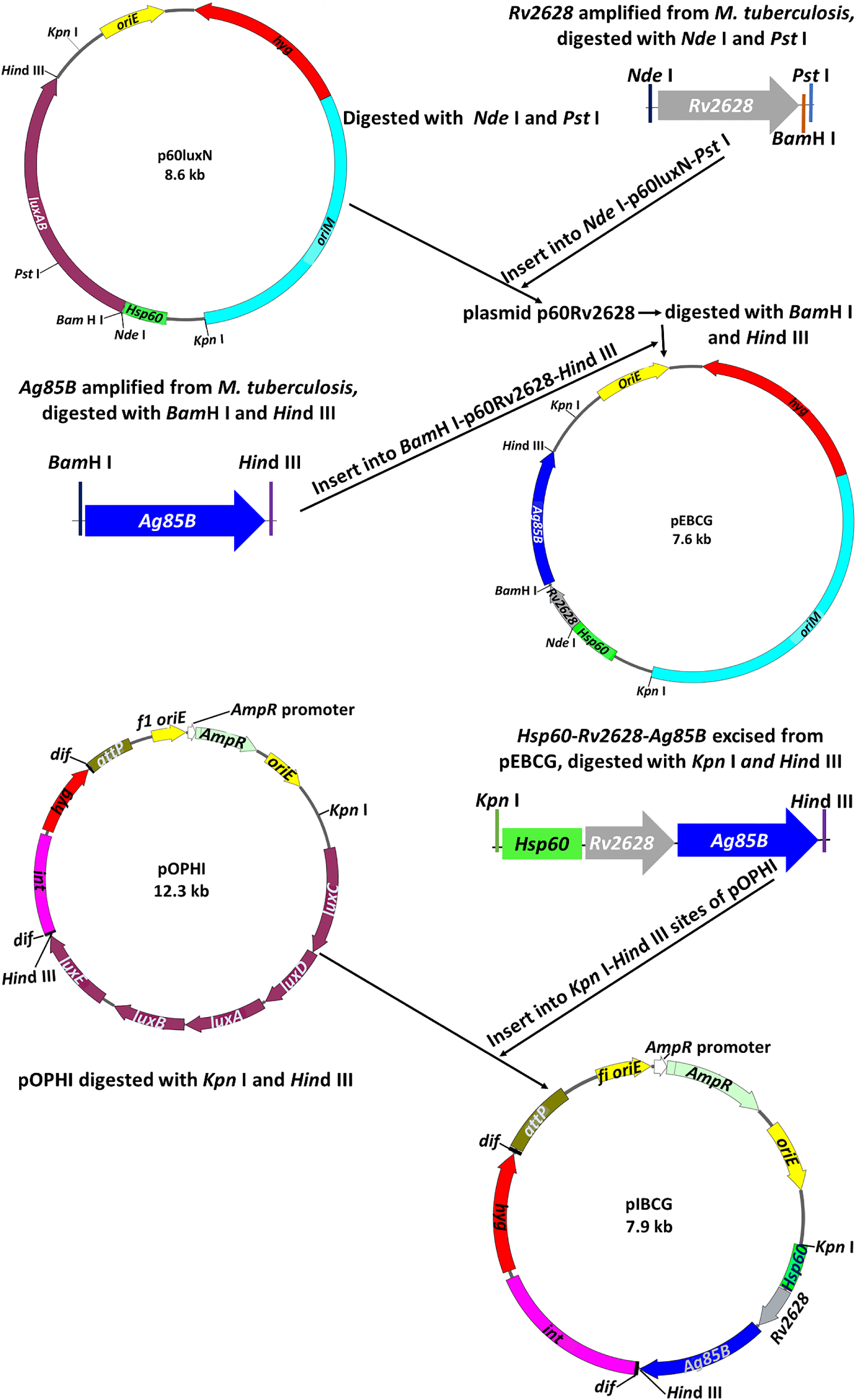
The schematic presentation for constructing the plasmids used for engineering the recombinant DR-BCGs. The Rv2628 gene was inserted next to the hsp60 promoter on the *Nde* I/*Pst* I site followed by Ag85B on the *Bam*H I/ *Hin*d III sites of plasmid p60LuxN to create the extrachromosomal plasmid pEBCG. The *hsp60-Rv2628-Ag85B* cassette from the pEBCG plasmid was thereafter excised and inserted on the *Kpn* I/ *Hin*d III sites of the integrative vector pOPHI to create the integrative plasmid pIBCG. The two plasmids were transformed into the competent cells of drBCG to create the recombinant drug-resistant BCG-E and RdrBCG-I respectively. **Abbreviations**: *attP*, mycobacteriophage L5 attachment site; *dif*, deletion-induced filamentation of *E. coli*; *int*, integrase gene; *AmpR*, the ampicillin resistance gene; *oriE*, the origin of replication of *E. coli*; *oriM*, thermosensitive origin region of mycobacteria; *hyg*, hygromycin resistance gene; hsp60, mycobacterial hsp60 promoter.

We therefore transformed by electroporation the S-L-E-P-A-resistant BCG mutant (designated as “drBCG”) with either plasmid pEBCG or pIBCG (Fig. 1). Interestingly, PCR amplification of the transformants showed the expected band corresponding to the 0.6-kb fragment spanning the *Rv2628* and *Ag85B* genes. Additionally, sequencing of the transformants obtained on hygromycin-containing plates resulted in sequences authentic to the design. This suggests that transformation was successful and that the colonies were RdrBCG-E and RdrBCG-Ihi.

### Exogenous genes stably existed in RdrBCG-I both *in vitro* and *in vivo*

To determine if the RdrBCG strains can maintain the extra *Ag85B* and *Rv2628* in the absence of selective pressure as would be the case in vaccinated mice, we assessed the presence of the two exogenous genes by PCR after five passages in liquid 7H9 broth. The RdrBCG-Ihi maintained a steady presence of the incorporated extra *Ag85B* and *Rv2628* genes after 5 passages (about 45 generations) from construction (Fig. S1B). However, the *hyg* and *int* genes were lost in all the 50 randomly selected colonies designated as RdrBCG-I. The fragment *dif-hyg-int-dif* was possibly excised by the endogenous XerCD recombinases of BCG leaving a *dif* in the genome. Likely, the removal of *int* gene could make the other integrated genes in the BCG genome more stable, as the integrase encoded by *int* cannot only integrate the plasmid containing *attP* site into the genome of BCG but also dissociate it at a low frequency (32). On the contrary, the exogenous genes in RdrBCG-E were lost after the fourth and fifth passage cycles in liquid 7H9 broth (Fig. S1B). This suggests that the pEBCG plasmid was not stable which prompted us to only use the RdrBCG-I for further studies. Similarly, the fifty RdrBCG-I colonies randomly isolated from lungs/spleens of 5 SCID mice 5 months after challenge in the virulence study below, maintained a steady presence of the exogenous *Ag85B* and *Rv2628* (Fig. S1C). This showed that the fragment *hsp60*-*Rv2628-Ag85B* existed very stably in RdrBCG-I both *in vitro* (100% about 45 generations) and *in vivo* (100%, ≥ 20 weeks) in the absence of the selection antibiotic.

### Rifampin-resistant *M. tuberculosis* (RR-*M. tuberculosis*) isolated

Considering that rifampin is the most important drug of modern TB chemotherapy as it has unique bactericidal and sterilizing activity against *M. tuberculosis* and that rifampin-resistant TB is base of all forms of serious DR-TB, we thought it necessary to use a RR-*M. tuberculosis* strain in our vaccine efficacy study. Furthermore, using RR-*M. tuberculosis* would make it easy to distinguish *M. tuberculosis* from the RdrBCG which cannot grow on rifampin containing plates for it was sensitive to rifampin. Upon screening the *M. tuberculosis* H37Rv, we obtained 3 spontaneous mutants on 7H11 plates containing rifampin 16 µg mL^−1^. Compared to the wildtype *M. tuberculosis* H37Rv, we noted an increased MIC for rifampin (≧ 64 µg mL^−1^) against the three isolated mutants. This suggested that the isolates were truly resistant to rifampin. Genotypic resistance was confirmed by PCR amplification of the RR-*M. tuberculosis* colonies using the rpoBf/rpoBr primer pair (Table 3) and sequencing. We obtained a spontaneous SNP at codon 531-TCG to TTG (Ser→Leu) previously reported to be associated with rifampin resistance (33, 34).

### DST of RdrBCG-I, WtBCG and RR-*M. tuberculosis*

We then tested the MICs of RdrBCG-I, WtBCG and RR-*M. tuberculosis* against selected anti-TB drugs (ALZP) to reconfirm their susceptibility levels. As expected, the MICs to RdrBCG-I were much higher for the drugs on which they were selected on compared with the parent WtBCG or *M. tuberculosis* H37Rv strains (Table 2). Likewise, the rifampin MIC to the RR-*M. tuberculosis* was much higher than that of the *M. tuberculosis* H37Rv and BCGs’. These findings suggested that the RdrBCG-I could not likely be killed by ALZP when vaccinated to mice adjunct to such drugs in the murine model. On the other hand, the BCG’s susceptibility to rifampin would allow easy quantification of the RR-*M. tuberculosis* from the vaccinated mice cultured on rifampin containing plates. Similarly, the levofloxacin containing plates were used to quantify RdrBCG-I by killing the mixed *M. tuberculosis* H37Rv, which was verified first *in vitro* using the mixture of different ratios of RdrBCG-I to RR-*M. tuberculosis*.

**Table 2:** The MICs of selected anti-TB drugs against RdrBCG-I, WtBCG and *M. tuberculosis* H37Rv strains

| Drugs | Strains/ MICs ($\mu\text{g mL}^{-1}$) | | | |
| --- | --- | --- | --- | --- |
|  | RdrBCG-I | WtBCG | *M. tuberculosis* H37Rv | RR-*M. tuberculosis* |
| Streptomycin | 64 | 0.25 | 1 | 1 |
| Ethambutol | 5 | 0.5 | 0.5 | 0.5 |
| Amikacin | 5 | 0.5 | 0.5-1 | 0.5-1 |
| Levofloxacin | 4 | 0.5 | 0.5 | 0.5 |
| Prothionamide | 64 | 0.25 | 0.5 | 0.5 |
| Rifampin | 0.25-0.5 | 0.025 | 0.25 | $\geq 256$ |
| Pyrazinamide * | / | / | 100-150 | 100-150 |

\*We did not test the MIC of pyrazinamide to BCG strains as it is well known that BCG is naturally resistant to pyrazinamide. Here the MIC of pyrazinamide to *M. tuberculosis* strains was tested using the MGIT 960 (Becton, Dickinson and Company. USA).

**Table 3:**
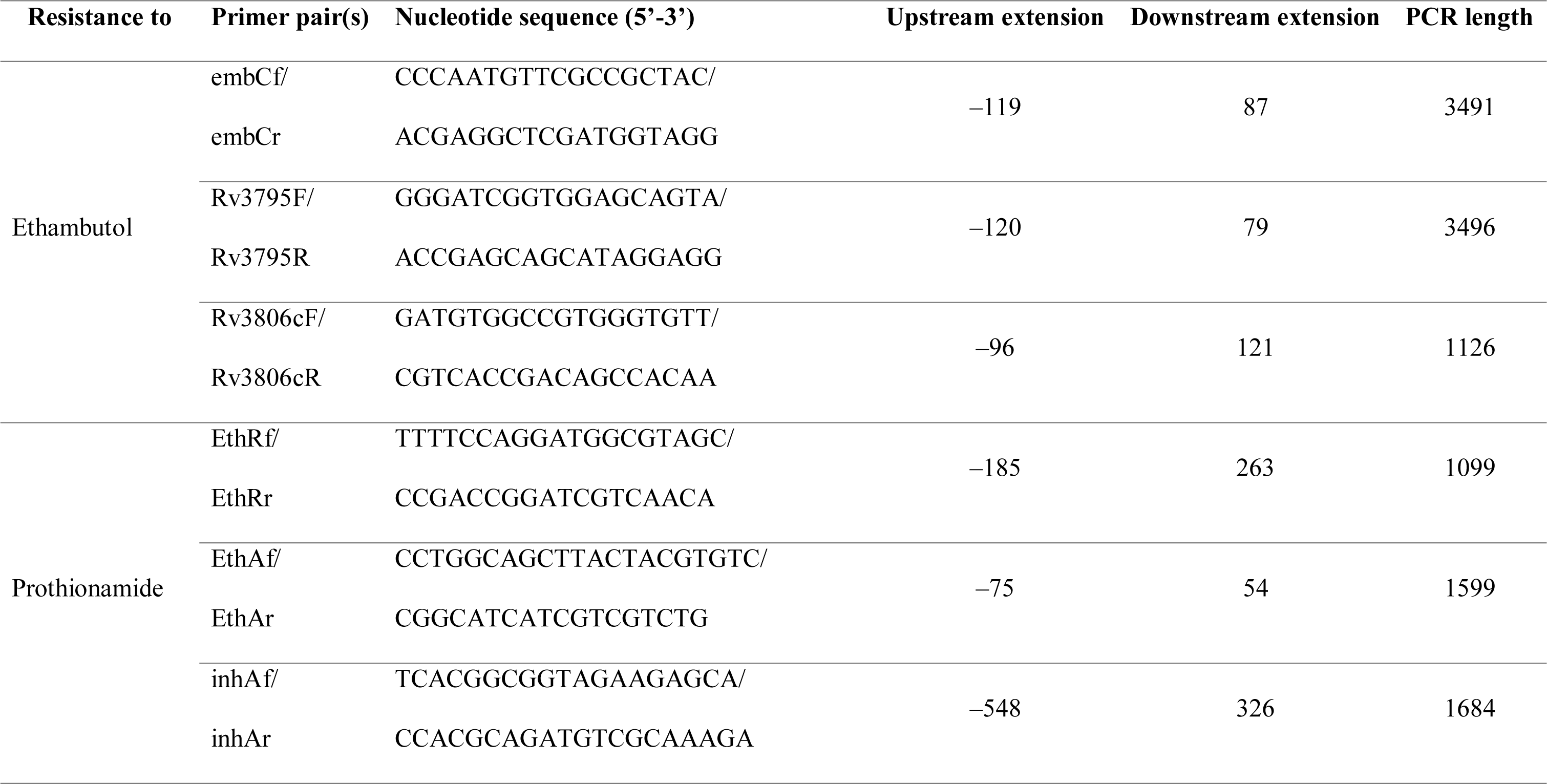
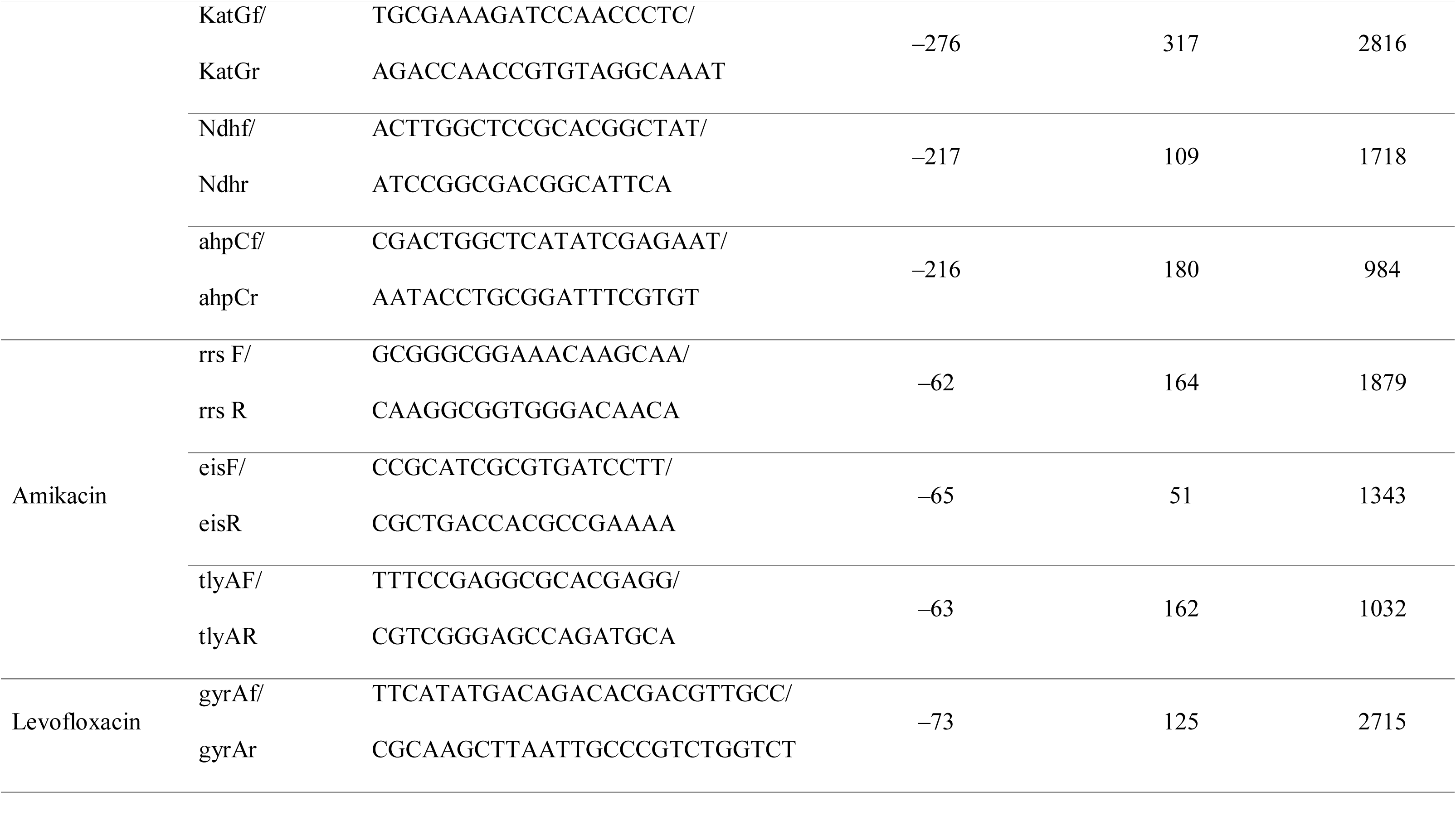
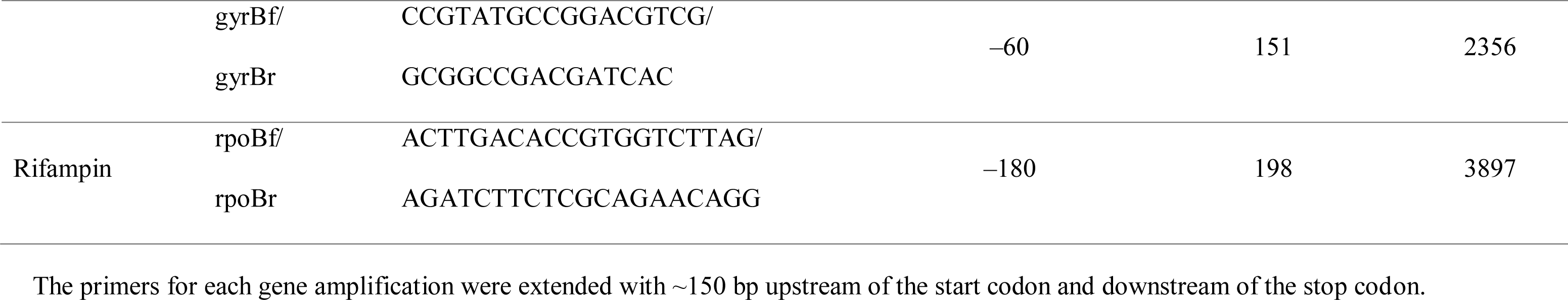
PCR and sequencing primers used to delineate target-based spontaneous genotypic resistance of the DR-BCGs and RR-*M. tuberculosis*

### RdrBCG-I inhibits the growth of *M. tuberculosis in vitro*

To test if the RdrBCG-I has an inhibitory effect on the growth of the pathogenic *M. tuberculosis in vitro,* the RdrBCG-I was co-cultured with the selectable marker-free autoluminescent *M. tuberculosis* reporter strain (UAlRv) (32) in liquid media. We assessed the UAlRv growth by monitoring the relative light unit (RLU) pattern over 20 days. Co-culturing the RdrBCG-I with UAlRv lowered the RLU by 1 log and 2 logs in the most and less concentrated UAlRv, respectively (Fig. 2A-B). The decline in UAlRv growth was more noticeable from 4 days post incubation whereby the co-culture trailed the control group in RLU counts. Notably, the RLU decline corresponded with the increase in RdrBCG-I concentration. On the other hand, co-culturing the UAlRv with the WtBCG resulted in less than 1 log decline in RLU mL^−1^ regardless of the concentration of the culture by the 20^th^ day post incubation (Fig. 2C-D). This suggested that the RdrBCG-I interfered with *M. tuberculosis* H37Rv growth more considerably than the WtBCG. So, we hypothesized that RdrBCG-I would complement chemotherapy in a DR-TB regimen against the *M. tuberculosis* for RdrBCG-I is resistant to the drugs used. We then co-cultured the RdrBCG-I and UAlRv in the presence of levofloxacin (L) and prothionamide (P) (drugs to which the RdrBCG-I is resistant). We only opted for L and P instead of ALZP due to pyrazinamide’s lack of activity *in vitro,* especially at normal pH. Additionally, we omitted amikacin as it was only used in the intensive phase before therapeutic vaccination in the vaccine efficacy study. Interestingly, we observed that the most concentrated UAlRv reached the baseline RLU level on 20^th^ day post incubation when co-cultured with RdrBCG-I while the baseline was reached in 7 days when the RdrBCG-I was cultured with a 10-fold diluted UAlRv supplemented with L and P (Fig. 2E-F). On the contrary, we did not achieve baseline RLU level when the WtBCG was co-cultured with the most concentrated UAlRv in the presence of L and P (Fig. 2G). We only obtained baseline RLU when the WtBCG was co-cultured with a 10-fold diluted UAlRv supplemented with L and P (Fig. 2H).

**Figure 2:**
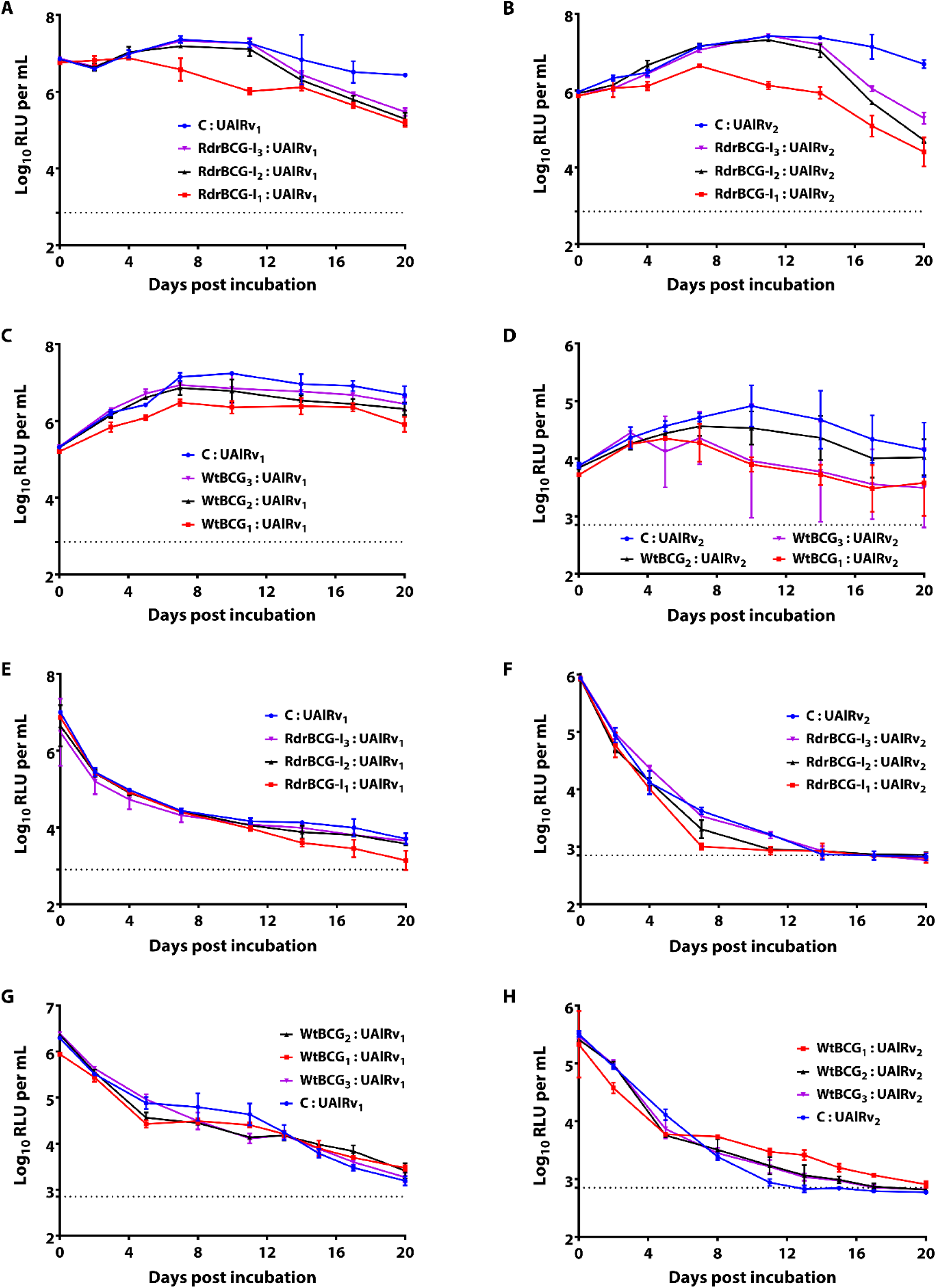
The RdrBCG-I inhibits *M. tuberculosis* growth *in vitro.* Co-culturing the RdrBCG-I and UAlRv lowered the UAlRv’s RLU count in a concentration-dependent manner and enhanced chemotherapy in lowering the RLU. Effect of RdrBCG-I (A-B) and WtBCG (C-D) on *M. tuberculosis* without drugs. Effect of RdrBCG-I (E-F) and WtBCG (G-H) on *M. tuberculosis* in the presence of levofloxacin and prothionamide. **Abbreviations:** C, control (no BCG); Subscripts 1, 2 and 3 denote undiluted, 10-fold dilution and 100-fold dilution for both BCG and UAlRv; the horizontal dotted lines denote the background log_10_ RLU mL^−1^. Results are mean ± SD of the log_10_RLU / 1 mL sample (n = 3 tubes per each dilution per sample). These experiments were performed in three parallel groups and repeated three times and only representative results are shown.

### Testing the safety and virulence of RdrBCG-I vaccine

The safety of RdrBCG-I was assessed by intravenously challenging 5-7 weeks old female severe combined immune deficient (SCID) mice with either the RdrBCG-I or WtBCG. A day after infection, the colony-forming units (CFU)/organ detected were 3.09 ± 0.03 log_10_ CFU/lung and 3.21 ± 0.04 log_10_ CFU/spleen of the WtBCG. On the other hand, 3.05 ± 0.08 log_10_ CFU/lung and 3.30 ± 0.02 log_10_ CFU/spleen of the RdrBCG-I were detected in the challenged mice (Fig. 3). Notably, there was no statistically significant difference in the number of detected BCG between the WtBCG and RdrBCG-I groups with a *P*-*value* of 0.6126 (lung) and 0.1055 (spleen). Seven days after inoculation, the pulmonary and splenic BCG loads increased to 4.00 ± 0.05 log_10_ CFU/lung and 4.30 ± 0.03 log_10_ CFU/spleen for the WtBCG challenged mice. On the other hand, the RdrBCG-I had grown to 3.90 ± 0.04 log_10_ CFU/lung and 4.20 ± 0.04 log_10_ CFU/spleen. Again, we observed no difference in the lung and splenic BCG loads (*P=* 0.3598 and 0.1093, respectively). Four weeks post challenge, 5.71 ± 0.00 vs 6.13 ± 0.03 log_10_ CFU were retained in the lungs of WtBCG and RdrBCG-I challenged SCID mice, respectively with a *P-value* of 0.0003. Additionally, 6.52 ± 0.02 vs 7.46 ± 0.04 log_10_ CFU/spleen were retained in WtBCG and RdrBCG-I challenged mice, respectively (*P*-*value* < 0.0001). As shown in Fig. 3A-B, the trend continued until the final day of the observation period (20 weeks).

**Figure 3:**
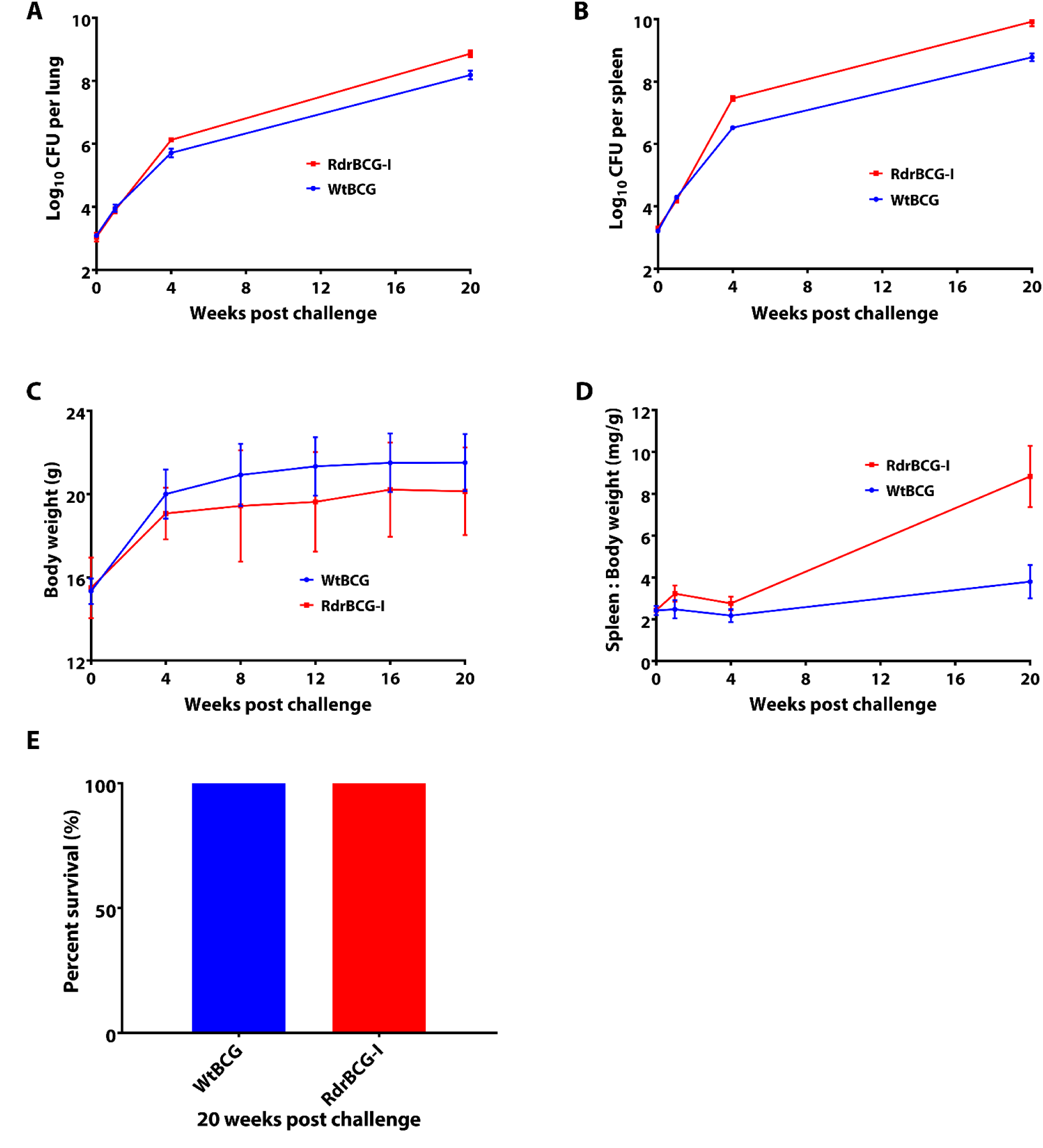
RdrBCG-I and WtBCG growth dynamics in SCID mice plus mice body and spleen weight changes as well as survival rates over time. More RdrBCG-I were retained in the lungs **(A)** and spleens **(B)** than WtBCG with a *P-value* of 0.0001. Additionally, RdrBCG-I challenged mice had lower body weights **(C)** but bigger spleen to body weight ratios **(D)** than WtBCG challenged mice (*P* < 0.001). Mice survival rates **(E)** in both BCG types. Error bars = mean ± S.D for the Log_10_ CFU/organ, body weights as well as spleen weight/body weight ratio

From 4 weeks post challenge, we noted that SCID mice challenged with RdrBCG-I had lower body weight (Fig. 3C) but had an overall bigger spleen to body weight ratio (Fig. 3D) than their WtBCG challenged counterparts (*P <* 0.001). Most importantly, 20 weeks after challenge, all the SCID mice were still alive (100% survival rate) regardless of the BCG strain they were inoculated with (Fig. 3E), which showed that the two BCGs (RdrBCG-I and WtBCG) were very safe.

### RdrBCG-I increased the therapeutic efficacy of a second-line anti-TB regimen in a RR-TB murine model

To evaluate the RdrBCG-I’s efficacy as a therapeutic vaccine, RR-*M. tuberculosis*-infected mice were first treated with the regimen; amikacin (A), levofloxacin (L), pyrazinamide (Z) and prothionamide (P) for 2 months (2ALZP) and later followed by 3 months administration of L, Z and P (3LZP). In addition to the drugs, the mice were vaccinated with three doses of RdrBCG-I or WtBCG administered at the end of month 2, 3, and 4 of treatment by either the aerosol (a.s) or subcutaneous (s.c) route (Table 4). The detailed lung CFU counts recovered at various time points are presented in Table S1. The day after infection (D -15), 2.14 ± 0.07 log_10_ CFU/lung was detected. At the time of treatment initiation (Day 0), the RR-*M. tuberculosis* burden had risen to 6.39 ± 0.1 log_10_ CFU/lung suggesting that the RR-*M. tuberculosis* was favorably multiplying in the mice. One month after treatment initiation, all five untreated mice were very sick and were sacrificed. At the time of their sacrifice, the RR-*M. tuberculosis* log_10_ CFU/lung in the untreated group reached 8.05 ± 0.35, which indicated that the RR-*M. tuberculosis* we got was virulent. At the end of two months of treatment with ALZP, the RR-*M. tuberculosis* bacterial burden declined from 6.39 ± 0.10 log_10_ CFU/lung as a baseline to 4.48 ± 0.14 log_10_ CFU/lung. At the beginning of month 3 of treatment, mice were switched to LZP with or without BCG vaccination as shown in Table 4. The detailed BCG doses administered at each time point for both the aerosol and subcutaneous route is given in Table S2 while the amount of RdrBCG-I recovered in the lungs of vaccinated mice is given in Table S3 as well as Fig. S2. For the aerosol route, the vaccination doses (log_10_ CFU/lung) of WtBCG/RdrBCG-I were 1.95 ± 0.04/2.00 ± 0.05 for the prime dose; 2.02 ± 0.06/1.99 ± 0.04 for the first booster dose, and 1.98 ± 0.06/2.02 ± 0.07 for the second and final booster dose respectively. On the other hand, the vaccination doses for the subcutaneous route ranged from 2.0 to 6.0 × 10^5^ CFUs per jab. As shown in Fig. 4A, chemotherapy significantly lowered the RR-*M. tuberculosis* bacterial load from 6.39 ± 0.10 log_10_ CFU lung at the time of treatment initiation to 4.48 ± 0.14 log_10_ CFU/lung after two months of chemotherapy (*P* < 0.0001) and remained almost constant thereafter.

**Figure 4:**
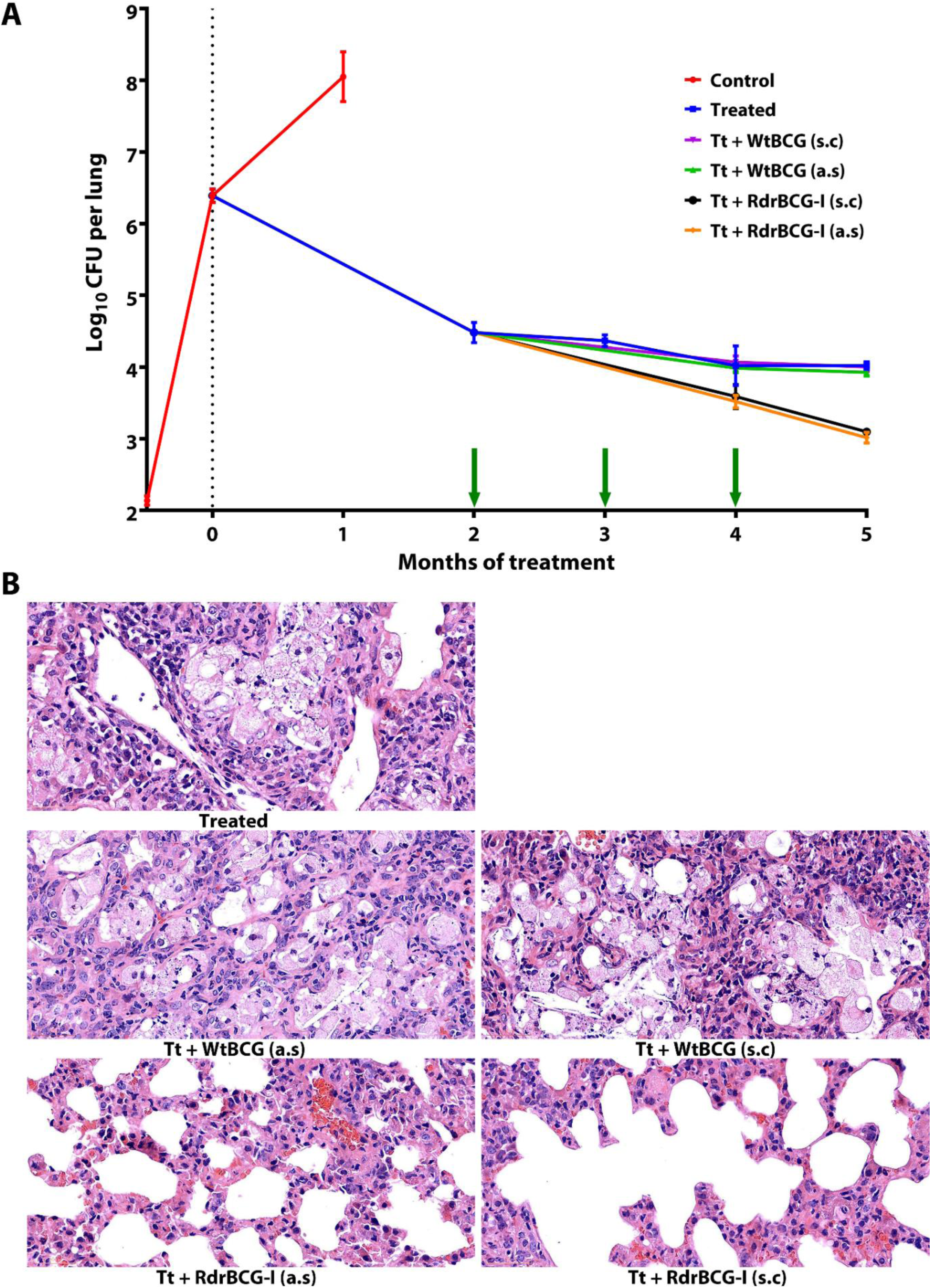
The effect of RdrBCG-I when added to a second-line drug regimen in a mouse model of DR-TB. **(A)** CFU counts during the experiment showing that RdrBCG-I complements chemotherapy and enhance *M. tuberculosis* clearance in infected mice lungs. Data (mean log_10_ CFU) were obtained from the lungs of five mice per time point, and the error bars represent the SD. The green arrows indicate the vaccination time points. **(B)** Histopathology of mice lungs from different groups at the end of 5 months of treatment (magnification: × 400). Treated (Tt) group manifested accumulation of multinucleated giant cells and caseous necrosis with alveolar foamy macrophages and necrotic acute inflammatory exudates in the alveolar spaces. Tt + WtBCG (a.s) and Tt + WtBCG (s.c) groups manifested accumulation of acute inflammatory exudates and dead macrophages with tissue debris plus alveolar macrophages with necrotic acute inflammatory exudates in the alveolar spaces. Tt + RdrBCG-I (a.s) and Tt + RdrBCG-I (s.c) groups manifested remodeling of the alveolar septae with fibrous connective tissues as well as atrophy of the alveolar wall. **Abbreviations**: Tt, treated with 2ALPZ/3LPZ; A, amikacin (100 mg kg^−1^); L, levofloxacin (300 mg kg^−1^), P, prothionamide (25 mg kg^−1^); Z, Pyrazinamide, (150 mg kg^−1^); a.s, aerosol vaccination; s.c, subcutaneous vaccination; 2ALPZ/3LPZ, 2 months of ALPZ followed by 3 months of LPZ; Control mice did not receive any treatment.

**Table 4:** Experimental scheme to test the efficacy of the RdrBCG-I as a therapeutic vaccine in a murine model of DR-TB

| Group name | Number of mice used to determine lung CFU counts per time point |  |  |  |  |  |  |  |
| --- | --- | --- | --- | --- | --- | --- | --- | --- |
|  | D -15 | D0 | M 1 | M 2 | M 3 | M 4 | M 5 | Total |
| Control | 5 | 5 | 5 |  |  |  |  | 15 |
| Treated (Tt) |  |  |  | 5 | 5 | 5 | 5 | 20 |
| Tt + WtBCG (a.s) |  |  |  | V | V | V 5 | 5 | 10 |
| Tt + WtBCG (s.c) |  |  |  | V | V | V 5 | 5 | 10 |
| Tt + RdrBCG-I (a.s) |  |  |  | V | V | V 5 | 5 | 10 |
| Tt + RdrBCG-I (s.c) |  |  |  | V | V | V 5 | 5 | 10 |
| Total | 5 | 5 | 5 | 5 | 5 | 25 | 25 | 75 |

Control, untreated group; Treated (Tt), only treated with 2ALPZ/3LPZ= 2 months of ALPZ followed by 3 months of LPZ; a.s, aerosol vaccination; s.c, subcutaneous vaccination; D, day; D -15, the day after infection; D0, day of treatment initiation; M1, one month after treatment initiation and so on; V, vaccination. Drug dosage (mg kg<sup>-1</sup>): A, amikacin (100); L, levofloxacin (300); P, prothionamide (25); Z, pyrazinamide (150). Mice were sacrificed 3 days after administration of the last treatment dose.

Compared with the treated only group, additional vaccination with RdrBCG-I by the aerosol route showed lower RR-*M. tuberculosis* loads (log_10_ CFU/lung): 4.02 ± 0.28 (treated) vs 3.52 ± 0.09 (Tt + RdrBCG-I (a.s), *P* = 0.0049) and 4.02 ± 0.05 (treated) vs 3.01 ± 0.08 (Tt + RdrBCG-I (a.s), *P* < 0.0001) at the end of 4 months and 5 months of treatment respectively (Fig. 4A). Likewise, compared with the treated only group, additional vaccination with RdrBCG-I by the subcutaneous route also showed lower RR-*M. tuberculosis* loads (log_10_ CFU/lung): 4.02 ± 0.28 (treated) vs 3.59 ± 0.17 (Tt + RdrBCG-I (s.c), *P* = 0.0179) and 4.02 ± 0.05 (treated) vs 3.10 ± 0.04 (Tt + RdrBCG-I (s.c), *P* < 0.0001) at the end of 4 months and 5 months of treatment respectively (Fig. 4A). The RR-*M. tuberculosis* burdens from different RdrBCG-I vaccination routes did not differ at all time points, suggesting that the route of vaccination might not have influenced the efficacy of the RdrBCG-I. Vaccination with WtBCG by either route did not significantly lower RR-*M. tuberculosis* burden after 2 vaccination doses (CFUs counted after 4 months of treatment, Fig. 4A) with a *P-value* of 0.7945 and 0.7091 for the aerosol and subcutaneous route, respectively. On the other hand, vaccination with WtBCG by aerosol route only lowered the RR-*M. tuberculosis* burden slightly after 3 doses (CFUs counted after 5 months of treatment, Fig. 4A, *P* = 0.0218) while the subcutaneous route did not affect the burden significantly (*P* = 0.6329). Most importantly, vaccination with RdrBCG-I continuously lowered the RR-*M. tuberculosis* load unlike in the treated control and WtBCG vaccinated groups whose RR-*M. tuberculosis* loads stagnated at around 4 log_10_ CFU after two months of drug therapy.

Since the RdrBCG-I is intended for use as a therapeutic vaccine to improve DR-TB treatment outcome, it was essential to evaluate if its administration could minimize pulmonary tissue pathology thereby improving the quality of life after treatment. We stained the mice lung tissues with hematoxylin and eosin (H&E) to monitor the morphological changes at the end of the study duration. Pathological examination revealed the presence of inflammatory cells and multiple foamy macrophages in the alveolar spaces of the mice lung sections stained with H&E. The inflammatory cells and foamy macrophages were so prominent among the treated control and WtBCG co-treated groups that they at times completely obliterated the spaces (Fig. 4B). In contrast, the RdrBCG-I co-treated animals had minimal to no presence of foamy macrophages and inflammatory cells which was suggestive of the RdrBCG-I’s ability to minimize tissue pathology. Additionally, the RdrBCG-I co-treated animals manifested remodeling of the alveolar septae with fibrous connective tissues as well as atrophy of the alveolar wall, a feature of healing indicating that RdrBCG-I facilitated lung tissue healing (Fig. 4B).

## Discussion

The rise of DR-TB has put a dent in the fight against TB with would-be new drugs being rendered useless even before they are rolled out. It is in this view that therapeutic vaccines are a preferred choice for eliminating the pathogen as they cannot be affected by the pathogen’s resistance to the administered drugs (35). In the present study, a therapeutic RdrBCG vaccine was created by incorporating the DNA sequences encoding *M. tuberculosis’ Rv2628* and *Ag85B* into the plasmids pEBCG and pIBCG which were later transformed into the competent cells of drBCG. The vaccine’s drug-resistance was targeted against anti-tuberculous drugs used in the treatment of DR-TB, especially amikacin, levofloxacin, prothionamide as well as streptomycin. The vaccine’s stability was assessed by PCR amplification of a fragment spanning the inserted genes after serial passages *in vitro* and *in vivo*. We found that the RdrBCG transformed with pIBCG retained the exogenous genes about 50 generations after the initial transformation *in vitro* and in mice (Fig. S1B-C) for a long period implying that the RdrBCG-I vaccine can sustain its antigenic properties during the continuation phase of TB treatment.

Safety is one of the key attributes of a good vaccine. We used SCID mice which lacks both T and B lymphocytes as well as immunoglobulins to evaluate the safety of WtBCG and the RdrBCG-I (36). Although the inoculum sizes were similar in the two groups, RdrBCG-I challenged SCID mice had more BCG retained in both lungs and spleens than WtBCG challenged mice by 4 weeks post infection (Fig. 3A and 3B). In both BCG groups, no mouse died, which suggested that the vaccine was safe to be administered to mice. However, as shown in Fig. 3C and 3D, the RdrBCG-I challenged mice had overall low body weight but bigger spleen to body weight ratios than their wild-type counterparts. Since big spleen size can be a sign of immunogenicity (37), it would, therefore, imply that RdrBCG-I elicits immune reaction more than the WtBCG. All these could probably be because of the inserted genes which may have enhanced the RdrBCG’s growth and replication *in vivo*.

Competition studies between bacteria have been used to delineate their interactions both *in vitro* and *in vivo* thereby being used to explore the potential mechanism of action in health and disease (38). For instance, bacteria may compete for resources, binding sites, or in some cases excrete substances unfavorable to their competitors which may limit the growth or survival of other bacteria (39). Herein, co-culturing the RdrBCG-I and the pathogenic *M. tuberculosis in vitro,* yielded minimal UAlRv growth as observed by declining RLU counts beginning day 4 post incubation. Most importantly, the most concentrated RdrBCG-I expedited the RLU decline which showed that the competitive effect of RdrBCG-I was titer or concentration-dependent. Considering that BCG and *M. tuberculosis* share homology (40), it is unlikely that BCG excreted substances that impacted UAlRv growth. As such, the plausible explanation is resource consumption whereby the RdrBCG-I may have outcompeted the *M. tuberculosis*. Furthermore, our findings suggest that the RdrBCG-I complements chemotherapy as it significantly lowered the duration required to kill *M. tuberculosis in vitro* in the presence of levofloxacin and prothionamide.

The RR-*M. tuberculosis*-infected BALB/c mice were treated with amikacin, levofloxacin, pyrazinamide and prothionamide for 2 months (intensive phase) and later followed by 3 months (continuation phase) administration of levofloxacin, pyrazinamide and prothionamide (2ALPZ/3LPZ) as shown in Table 4. We noted that drug treatment significantly lowered lung RR-*M. tuberculosis* load by 2 log_10_ CFU/lung after two months and remained almost constant thereafter (Fig. 4A) which is similar to previously reported findings by Grosset et al (41) as well as our previous study (42). This is likely due to the bacteria being in a persistent state which necessitates the prolongation of treatment in DR-TB (43). It is well noted that rifampin is the key drug in the current short-course TB treatment, such that when it is missing in a regimen or if a patient gets infected with a RR-*M. tuberculosis*, it is very hard to cure that patient, as rifampin can continuously kill the persisters uniquely (41). Interestingly, co-treatment with the RdrBCG-I significantly lowered the RR-*M. tuberculosis* burden during the continuation phase of treatment in the absence of rifampin (*P* < 0.0001). Most importantly, the decline was steady and continuous which would allow its administration during the subsequent months whilst on drug treatment. A recent report on clinical isolates showed that > 90% of isoniazid-resistant *M. tuberculosis* had cross-resistance with prothionamide as they have one same target (44). Such that a prothionamide containing regimen might not be as effective in such a scenario, but our findings showed that the RdrBCG-I could be of utmost help.

Studies have shown that *Ag85B* and *Rv2628* elicit Th1 immune response which is capable of killing *M. tuberculosis* (45, 46). We can, therefore, hypothesize that the prime RdrBCG-I and its booster doses allowed the upregulation of the Th1 response which resulted in the continuous killing of the non-replicating RR-*M. tuberculosis*. This phenomenon was, however, not observed in the WtBCG vaccinated groups (Fig. 4A) likely due to its susceptibility to the administered drugs which may have killed or strongly inhibited it before it even elicited any immune response (13, 47). Considering that the treatment of DR-TB is long which sometimes leads to treatment interruption in the affected patients (4), it will be beneficial to try out therapeutic vaccination strategy using vaccines like RdrBCG-I for it has shown the potential of significantly lowering the RR-*M. tuberculosis* burden in mice continuously during the continuation phase. Further studies using longer chemotherapy with/without RdrBCG-I and with relapse rates as the final index of therapeutic evaluation need to be carried out in the future. It should be noted that though we just used 2ALPZ/3LPZ in our regimen, S may replace A, as both streptomycin and amikacin are injectable drugs to which some clinical DR-*M. tuberculosis* isolates showed no cross-resistance. In our case, we just used amikacin for 2 months, but amikacin is recommended to be used for 6 months in patients with very good activity and low price. We can therefore speculate that if amikacin were given for one more month together with RdrBCG-I, the bactericidal activity of the Tt + RdrBCG groups could have been even better. In addition, according to the clinical DST results or even without the DST results, ethambutol can be added to this regimen as the resistance rate of clinical *M. tuberculosis* isolates to the first-line anti-TB drug ethambutol is relatively low. Most importantly, ethambutol is capable of preventing the emergence of drug resistance even in immunosuppressed state (48).

Previous studies on vaccination routes demonstrated that the aerosol route is more efficient in enhancing pulmonary *M. tuberculosis* specific cell mediated immune response than the subcutaneous route (49, 50). Essentially, the resultant cell mediated immune response, thus, CD4^+^, CD8^+^ T cells and related Th1 cytokines necessitate the pathogen’s clearance from the host (51). Similarly, this study found that aerosol vaccination of both the WtBCG and RdrBCG-I led to lower RR-*M. tuberculosis* burden in the lungs when compared to the subcutaneous route although the difference was not statistically significant (Tt + WtBCG (a.s) vs Tt + WtBCG (s.c) *P* values were 0.1202 and 0.0689 at 4 and 5 months respectively, while for Tt + RdrBCG-I (a.s) vs Tt + RdrBCG-I (s.c) *P* values were 0.437 and 0.0581 at 4 and 5 months respectively).

A good treatment outcome is not only marked by clearance of the infecting bacteria alone but the return of quality of life to pre-infection levels which help reduce disease sequelae in TB patients (52). To study the efficacy of the RdrBCG-I on improving the treatment outcome in DR-TB, lung pathological changes were assessed using H&E stain at the end of the 5^th^ month of treatment. Histopathological analysis of the lung tissues revealed the presence of numerous inflammatory cells spread all over the lung in the treated control and WtBCG co-treated groups (Fig. 4B). Since the presence of a pathogen leads to persistent inflammation (53), it would, therefore, mean that the RR-*M. tuberculosis* persisted more in the control and WtBCG vaccinated mice than RdrBCG-I co-treated animals. Furthermore, after five months of treatment, remodeling of the alveolar septae had already begun in the RdrBCG-I co-treated animals (Fig. 4B). It is worth noting that remodeling of the alveolar septae and fibrosis, leads to resolution (54, 55). Hence, we can speculate that RdrBCG-I facilitated lung tissue healing. Additionally, there was a tremendous presence of foamy macrophages and foreign body giant cells in the lung alveolar spaces of the treated control animals and those complemented with WtBCG than in the RdrBCG-I co-treated mice (Fig. 4B). Studies have shown that foamy macrophages provide an environment and nutrients for *M. tuberculosis’* survival (56). Furthermore, foamy macrophages inhibit the phagocytic and the host’s bactericidal activities thereby aiding the pathogen’s survival (56, 57). It is therefore not surprising that the treated control and WtBCG co-treated groups had more RR-*M. tuberculosis* retained in their lungs throughout the study duration than the RdrBCG-I vaccinated groups. Although aerosol vaccination may cause some inflammation in the lungs, severe reactions are usually during the early months when the *M. tuberculosis* loads are still very high. In our case, we administered the vaccine when *M. tuberculosis* loads had plateaued which is the recommended time point for therapeutic vaccination as it facilitates the killing of non-replicating bacteria (58). Our H&E findings also showed that the aerosol vaccination did not cause serious inflammation as there was no difference in pathological changes between the aerosol and subcutaneous vaccinated mice within the vaccine groups (Fig. 4B).

Finally, we acknowledge the concerns emanating from using the RdrBCG-I like potential dissemination of the drug resistance genes essentially creating superbugs. We, however, believe that the resistance genes in RdrBCG-I cannot be disseminated easily into *M. tuberculosis* in TB patients because all the resistance genes are in the genome of RdrBCG-I and not in the extra chromosome plasmid which could be transferred into other mycobacteria. Secondly, the *M. tuberculosis* cell wall composition is too complex which makes it very hard to transform the plasmid DNA in a mycobacterium into another mycobacterium. Therefore, we consider it an impossibility for the RdrBCG-I genomic DNA to be transferred into *M. tuberculosis* in TB patients and still be functional. Thirdly, we do not know any report about the spontaneous transfer of genomic DNA of one mycobacterium to another. Lastly, we believe that the mutations in some resistance genes like the *pncA* which encodes pyrazinamide activation enzyme, do not alter the susceptibility of *M. tuberculosis*. Even if the *M. tuberculosis* gained the resistance gene, it stills contains the wild-type gene which can still activate the prodrug making it susceptible. Another concern about the RdrBCG-I is whether it could be controlled (killed efficiently) by drugs in vaccinated patients or not. It should be noted that the RdrBCG-I is still sensitive to several anti-TB drugs, like rifampin, isoniazid and many other powerful drugs, such as linezolid and/or clofazimine. Furthermore, the RdrBCG is not virulent, hence there is a means of clearing it when the need arises. As the world continues searching for a lasting solution to combating DR-TB, we herein have demonstrated a promising potential therapeutic application of the RdrBCG-I against DR-TB.

Specifically, the RdrBCG-I complemented chemotherapy in lowering the lung RR-*M. tuberculosis* burden as well as minimized lung tissue pathology in vaccinated mice. In addition, RdrBCG-I persistence after cure of DR-TB could help patients from reinfection or disease relapse which is very common among previously treated patients. To the best of our knowledge, this is the first study to utilize a recombinant DR-BCG which allowed its co-administration with an anti-TB regimen to kill DR-*M. tuberculosis* significantly better. Such that going forward, using the “recombinant” plus “drug-resistant” BCG strategy could be a useful concept for developing therapeutic vaccines against DR-TB.

## Materials and Methods

### Screening and verification of the DR-BCG strains

To create the Rdr-BCG strain, it is essential to have a BCG strain which is resistant to the drugs used to treat TB as a base first. Evidence shows that different BCG strains have distinct but broad immunologic responses with no known front runner inducer of protective immunity in humans (59). To obtain a BCG mutant resistant to most of the commonly used anti-DR-TB drugs including streptomycin (S), levofloxacin (L), ethambutol (E), prothionamide (P) or amikacin (A), the BCG Tice which was proved to be among the top BCGs inducing strong protection (60) was used as a starter to get a series of DR-BCG mutants through screening on drug-containing plates. Briefly, the WtBCG (OD_600_: 0.8-1.0) was screened on Middlebrook 7H11 (Becton, Dickinson and Company, USA) agar plates containing different concentrations of selected anti-TB drugs. The plates were supplemented with 10% oleic-acid-albumin-dextrose-catalase (OADC) enrichment medium and drugs at concentrations indicated in Table 1. The plates were incubated at 37 °C for 4 to 5 weeks until evaluation. Since BCG is intrinsically resistant to pyrazinamide due to a natural mutation in its *pncA* gene which encodes the PZase for activating the prodrug pyrazinamide to its active form (61), it was not necessary to select pyrazinamide-resistant BCG. To obtain the target drBCG (BCG mutant resistant to S-L-E-P-A), we screened for spontaneous DR-mutants using different drugs and routes in parallel. For example, we used all the individual drug (S, L, E, P or A) containing plates to screen in the first run and verify the corresponding DR-BCG mutants. Then we started the second screening run based on the true DR-BCG mutants obtained and then start the follow-up runs until we got the true drBCG.

The isolated mutant colonies were cultured to mid-log phase (OD_600_ = 0.8 to 1.0) in Middlebrook 7H9 broth (Becton, Dickinson and Company, USA) supplemented with 10% OADC enrichment, 0.2% glycerol as well as 0.05% Tween 80. We then made tenfold serial dilutions of the DR-BCG liquid cultures and inoculated on 7H11 plates containing the same concentrations of the drug on which they were screened. This was done to verify if the mutants were truly resistant strains. In all instances, the WtBCG was plated as a control. Colonies were also amplified by PCR using various primer pairs for drug resistance (Table 3) to confirm if the obtained colonies were genotypically resistant to the drugs on which they were screened. The PCR products were sequenced and aligned with the reference sequence using the basic local alignment search tool (BLAST) (http://blast.ncbi.nlm.nih.gov/blast.cgi) to identify the mutation(s) associated with genotypic resistance.

### Construction of the RdrBCG strains

The schematic diagram of constructing the plasmids for creating the RdrBCG has been outlined in Fig. 1. Briefly, we amplified by PCR the fragments of 1-kb *Ag85B* and 0.3-kb *Rv2628* with the *M. tuberculosis* H37Rv genomic DNA as a template using Ag85Bf/Ag85Br and Rv2628f/Rv2628r primer pairs (Table S4). The PCR‒amplified DNA products were analyzed by electrophoresis in agarose gel and purified using PCR purification kits (Magen, China). PCR products were cloned into p60LuxN plasmid (62) in turn through classical cloning method. The purified *Rv2628* PCR product and plasmid p60LuxN were digested with *Nde* I/*Pst* I and then ligated and transformed into *E. coli* DH5α to obtain the plasmid p60Rv2628. Transformants were selected on hygromycin (200 µg mL^−1^) containing plates and were verified by PCR amplification with primers Rv2628f/Rv2628r and HygF/HygR, and sequencing (Table S4). The verified plasmid p60Rv2628 was digested with *Bam*H I/*Hin*d III and its backbone was ligated with the *Ag85B* PCR product digested with *Bam*H I/*Hin*d III. The ligation mixture was transformed into *E. coli* DH5α to create the plasmid pEBCG. Transformants were selected on hygromycin containing plates and verified by enzyme cut and then by PCR amplification with primers Ag85Bf/Ag85Br and HygF/HygR, and sequencing (Table S4). The sequencing results were all verified by alignment matching to the reference sequence using BLAST or manually. The plasmid pEBCG was a multicopy extra-chromosome plasmid in mycobacterium. Finally, we excised the *hsp60-Rv2628*-*Ag85B* cassette from plasmid pEBCG and subcloned it into the *Kpn* I/*Hin*d III sites of the integrative vector pOPHI (32) to create the integrative plasmid pIBCG. The pIBCG is a single copy plasmid whose *dif*-*int*-*hyg-dif* cassette containing the hygromycin resistance gene (*hyg*) and the integrase gene (*int*) can be removed by the endogenous XerCD recombinases of BCG (32).

Following plasmids pEBCG and pIBCG construction, we prepared competent cells of drBCG and transformed them by electroporation with verified plasmids pEBCG, pIBCG and p60luxNC (control) as previously described (63). Briefly, 40 mL of drBCG in Middlebrook 7H9 broth was centrifuged at 3000 RCF for 10 minutes and the pellet was washed three times in sterile ice-cold 10% glycerol. Thereafter, we added plasmid DNA (10 μL) and 200 μL of competent drBCG cells into 0.2 cm electroporation cuvette and incubated the mixture at 37 °C for 10 minutes. We followed this up by performing electroporation using a Bio-Rad Gene Pulser electroporation device (2.5 KV voltage, 1000 Ω resistance). The bacteria mixture was then washed off the electroporation cuvettes 2 times with 7H9 supplemented with OADC and transferred into 50 mL centrifuge tubes. We then incubated the bacteria mixture at 37 °C for 20 to 24 hours. The mixture was plated on hygromycin (50 μg mL^−1^) containing 7H11 agar plates supplemented with OADC and were incubated at 37 °C for 4 to 5 weeks. To test if the transformation was successful, we amplified by PCR the 0.6 kb fragment spanning the *Ag85B* and *Rv2628* region using MarkF/MarkR primer pair (Table S4). When positive, they were designated as “RdrBCG-E (containing plasmid pEBCG) or RdrBCG-Ihi (integrated with plasmid pIBCG with *hyg-int* genes), RdrBCG-C (containing p60LuxNC)”. Furthermore, we cultured the RdrBCG-Ihi in antibiotic-free broth for about 10 generations and then plated the diluted culture onto antibiotic-free plates to obtain single colonies. Thereafter, we counter-selected the RdrBCG-I (without *hyg-int* genes) colonies by selecting those that could only grow on antibiotic-free plates but not on hygromycin containing plates and verify them by PCR amplification and sequencing. At the same time, RdrBCG-E, RdrBCG-Ihi, RdrBCG-I and RdrBCG-C were stored at −80 °C for future use.

### *In vitro* and *in vivo* exogenous genes stability testing for the RdrBCG strains

To assess if the exogenously introduced *Ag85B* and *Rv2628* existed stably in the constructed RdrBCG-E and RdrBCG-I, we passaged (1:9 dilution) the BCGs five times in the absence of hygromycin with each passage achieving an OD_600_ of 1.0. After the 4^th^ and 5^th^ passage, the subcultures were serially diluted and plated on 7H11 with hygromycin only for RdrBCG-E and without hygromycin for both strains and incubated at 37 °C. Four to five weeks after incubation, the plates were inspected for growth, and random colonies were selected on hygromycin-free plates for PCR amplification of the 0.6-kb fragment spanning the *Ag85B* and *Rv2628* using the primer pair MarkF/MarkR (Table S4). Similarly, stability was also assessed on the RdrBCG-I colonies recovered from the lungs and spleens of SCID mice sacrificed at the end of the *in vivo* safety study (5 months after infection) by amplifying the 0.6-kb fragment spanning *Ag85B* and *Rv2628* using MarkF/MarkR primer pair as well. The *M. tuberculosis* H37Rv and WtBCG were used as negative controls.

### Preparation of the RR-*M. tuberculosis*

We opted for a RR-*M. tuberculosis* because it accounts for a majority of all DR-TB cases. Additionally, our RdrBCG was susceptible to rifampin hence it would be easy to count the *M. tuberculosis* colonies on rifampin containing plates because only the RR-*M. tuberculosis* would grow. The RR-*M. tuberculosis* strain used in this study was selected on rifampin containing 7H11 plates. Briefly, *M. tuberculosis* H37Rv colonies were grown in Middlebrook 7H9 supplemented with 10% OADC enrichment, 0.2% glycerol and 0.05% Tween 80 to mid-log phase. The culture was washed and concentrated to 1/10 of the original culture volume. We then plated the concentrated culture on 7H11 supplemented with OADC containing rifampin at concentrations (µg mL^−1^) of 4, 8, 16, 32 and 64. After four weeks of incubation at 37 °C, the plates were inspected for growth. To verify if the obtained colonies were truly resistant to rifampin, we selected individual colonies and re-cultured them in 7H9 + OADC + glycerol + Tween 80 and plated on 7H11 containing rifampin at the same concentrations used to screen the mutants. To determine if the obtained resistant colonies harbored a mutation in the *rpoB* gene, which is almost the only reported rifampin resistance causing gene, we amplified their genomic DNA using the rpoBf/rpoBr primer pair by PCR (Table 3). The PCR product was sequenced (BGI, China) and the obtained sequences were aligned to the reference sequence using BLAST to identify the spontaneous mutations.

### DST of the RdrBCG-I, WtBCG, *M. tuberculosis* H37Rv and RR-*M. tuberculosis*

We detected the resistance levels of RdrBCG-I to selected drugs including streptomycin, amikacin, levofloxacin, ethambutol, prothionamide and rifampin to ensure that it would not be killed if administered adjunct to chemotherapy in the RR-TB murine model. We aimed at confirming that the RdrBCG-I was sensitive to rifampin while the RR-*M. tuberculosis* was sensitive to the drugs selected in our regimen. To accomplish this, we tested the MICs of these drugs to RdrBCG-I and RR-*M. tuberculosis* with their parent strains as controls. The MICs were determined by the classical agar 7H11 media method as previously described with some modifications (64). Briefly, a series of 10-fold dilutions of 500 μL bacterial culture were separately plated on 7H11 solid media containing different 2-fold diluted concentrations of the drugs above. The MIC was defined as the lowest drug concentration which inhibited at least 99 % of the growth observed on the drug-containing plates compared to the drug-free control plates. Furthermore, we tested the MIC of pyrazinamide to *M. tuberculosis* strains using the MGIT 960 (Becton, Dickinson and Company. USA) per manufacturer’s protocol. As it is well known that BCG is intrinsically resistant to pyrazinamide (61), we did not test pyrazinamide susceptibilities of BCG strains.

Additionally, we tested which drug(s) could be used to differentiate the RdrBCG/WtBCG with RR-*M. tuberculosis* efficiently by plating different ratios of BCG/RR-*M. tuberculosis* on the plates containing 2-fold series of drugs selected. The standard was a drug concentration that could inhibit 99.9% growth of one species but would not affect the growth of the other bacteria. At last, rifampin 4 µg mL^−1^ and levofloxacin 2 µg mL^−1^ were selected to grow RR-*M. tuberculosis* and RdrBCG respectively. Unfortunately, we did not find a good drug/compound to grow WtBCG only while fully inhibiting *M. tuberculosis* in the mixture. Although cycloserine 30 µg mL^−1^ was reported to selectively inhibit the growth of *M. tuberculosis* and allow the growth of BCG (65), the high resistant rates among *M. tuberculosis* make it an unreliable drug for this purpose (66).

### *In vitro* growth curves of UAlRv co-cultured with WtBCG or RdrBCG-I

We then tested the *in vitro* growth kinetics of the UAlRv strain in the presence of RdrBCG-I or WtBCG in 7H9 liquid medium. The growth of UAlRv can be monitored in real-time by measuring RLU as the correlation between RLU and CFU is very high (67). Briefly, the UAlRv, WtBCG and RdrBCG-I were separately grown in 7H9 supplemented with tween 80 and OADC to mid-log phase. For the co-culturing experiments, the optical densities of the cultures were adjusted to the same value (OD_600_: 0.7). Thereafter, we prepared mixtures of the RdrBCG-I/UAlRv or WtBCG/UAlRv to a final volume of 200 µL as shown in Table S5. The culture mixtures were incubated at 37 °C and monitored for 3 weeks. Additionally, the culture mixtures were supplemented with levofloxacin 2.0 µg mL^−1^ as well as prothionamide 0.8 µg mL^−1^ to determine if the RdrBCG-I can complement the bactericidal activity of the two drugs. We did not use pyrazinamide as it does not have good activity *in vitro,* especially at normal pH. We also did not use amikacin for it was only used in the intensive phase in the TB murine model before vaccination with BCG. Control tubes containing UAlRv and 7H9 with or without drugs (levofloxacin and prothionamide) were included. Bacterial growth was monitored by checking the RLU (triplicates) using the Glomax 20/20 Luminometer on days 0, 2, 4, 7, 11, 14, 17, and 20 post incubation. These experiments were repeated three times even by different persons and only representative results have been shown.

### Testing the safety and virulence of RdrBCG-I

We tested the safety and virulence of the RdrBCG-I vaccine in SCID mice following the previous reports (68, 69) with WtBCG as a control. Briefly, sixty, 4-6 weeks old female SCID mice purchased from Charles River Laboratories (Beijing, China) were housed in a specific pathogen-free environment and had free access to food and water. Following a week of acclimatization, the mice were intravenously infected with either RdrBCG-I or WtBCG (OD_600_: 0.5) reconstituted in 200 µL PBS via the tail vein. Uninfected control mice were injected with the same volume of PBS. A day after challenge, 3 animals per BCG group were sacrificed and their lungs and spleens were homogenized and a series of 10-fold diluted homogenates were plated to quantify the bacterial inoculating dose. To analyze the growth of the BCG strains in the SCID mice, 5 mice from each BCG group were sacrificed at 1 week and 4 weeks after challenge. The CFUs were enumerated from both the lungs and spleens. The rest of the mice were monitored over time (20 weeks) to ascertain their survival/mortality rates. If no mouse died, they would be sacrificed at the end of 20 weeks after infection for detecting CFUs in the lungs and spleens. In addition, body weights were recorded once weekly while spleen weights were recorded on the day of sacrifice.

### Testing the therapeutic efficacy of RdrBCG-I

The therapeutic efficacy of the RdrBCG-I was tested in BALB/c mice challenged with the RR-*M. tuberculosis.* Briefly, 4 to 6 weeks old female BALB/c mice were purchased from Charles River Laboratories (Beijing, China) and housed under standard conditions of light and temperature. Food and water were provided ad libitum. Following a week of acclimatization, the animals were loaded onto a Glas-Col inhalation exposure system (Glas-Col, Terre Haute, IN. USA) with 15 mice per compartment. Thereafter, 10 mL of the RR-*M. tuberculosis* was loaded and blown for 40 minutes as per the system’s instructions. After infection, the animals were randomly divided into different groups of 5 mice/cage as outlined in Table 4. To determine the number of RR-*M. tuberculosis* successfully implanted in the lungs, five mice were sacrificed a day after infection (D-15) and their lungs were homogenized, diluted and plated. Also, we sacrificed five mice on the day of treatment initiation (D0) to determine the baseline pulmonary bacterial load.

Studies on immunization route(s) have shown that the aerosol or intranasal route of vaccination of BCG induces a stronger immune response than the subcutaneous or intradermal routes (49, 50). Furthermore, the aerosol route could be more tolerable as well as acceptable to TB patients as it has a relatively low risk for potential hypersensitivity reaction(s). As such, mice in this experiment were grouped to either the aerosol or subcutaneous routes with WtBCG as a control. For both BCG types, the OD_600_ was adjusted to 0.7 and the animals were vaccinated with three doses at monthly intervals during the continuation phase beginning the start of month 3 of chemotherapy. For the aerosol vaccinated mice, 10 mL of the BCG culture was administered via the Glas-Col inhalation exposure system (Glas-Col, Terre Haute, IN. USA). The system was calibrated as per the manufacturer’s protocol with an exposure time of 40 minutes. To determine the amount of successfully implanted BCG in the lungs for the aerogenically vaccinated mice, sentinel BALB/c mice were included at each run. The sentinel mice were sacrificed 24 hours after exposure and their lungs were cultured on 7H11 plates. For the subcutaneously vaccinated mice, 200 µL of each BCG was injected just under the skin of each mouse at the scheduled time as in Table 4. To quantitate the amount vaccinated, we cultured part of the stock BCG solution on 7H11. The plates were incubated at 37 °C for 4 to 5 weeks before evaluation.

To enumerate the RR-*M. tuberculosis* burden following drug therapy and vaccination, lungs were aseptically extracted and homogenized in 2 mL sterile PBS using glass homogenizers. Thereafter, tenfold serial dilutions of the homogenate were made and plated on 7H11 plates containing rifampin at 4 µg mL^−1^ to quantify RR-*M. tuberculosis* CFUs and on plates containing levofloxacin at 2 μg mL^−1^ to quantify RdrBCG-I CFUs. This was specifically done at months 3, 4, 5 after treatment initiation when the differentiation of the two bacteria was needed. At all-time points, plates were incubated at 37 °C for 4 to 5 weeks before counting the CFUs. Due to the lack of an effective selection method for the WtBCG, we did not quantitate its growth in the RR-*M. tuberculosis*-infected mice vaccinated with it.

To assess if the administered vaccine is effective in alleviating immunopathology in the lungs, we fixed lung sections in 10% neutral buffered formalin soon after euthanasia at the end of the 5^th^ month of treatment. The paraffin-embedded sections were cut and stained with H&E for histopathological examination by a pathologist who was blind to the experimental design.

### Drug(s) preparation and administration

All animals were treated with the same drug regimen from 15 days post infection till the final day of treatment. The drugs used were purchased from Dalian Meilun Biological Technology Co. Ltd, Dalian, China and were prepared weekly, reconstituted and administered as shown in Table S6. For administration, 200 µL of the drug solution containing the calculated drug concentration was drawn from the stocking container into a 1 mL syringe fitted with a gavage or an injection needle. The drawn solution was either given orally or subcutaneously (for amikacin) to each mouse for the designated duration as shown in Table S6. All drugs were given daily, five days per week.

## Ethical approval

All animal procedures followed the institutional guidelines on animal handling and were approved by the Institutional Animal Care and Use Committee of the Guangzhou Institutes of Biomedicine and Health (#2018041; #2018053).

## Statistical analysis

All experiments were performed in replicates where possible. The mean CFU/organ and RLU mL^−1^ were transformed to log_10_ before analysis. The log_10_ transformed data plus the mean mice weights for the different treatment and vaccination groups were then analyzed by T-test when comparing two groups and one-way analysis of variance (ANOVA) when comparing more than two groups. The GraphPad Prism 6 software (GraphPad, San Diego, CA) was used for all statistical analyses. The results were considered significant when *P* was < 0.05.

## Acknowledgments

We wish to thank the staff at the Guangzhou Institutes of Biomedicine and Health Pathology lab for processing the histopathology slides. Special thanks should also go to Mr. Erjun Zhang at the animal center who assisted us with animal handling during the vaccine safety study.

## Data and materials availability

All data associated with this study are present within the article and in its supplementary materials.

## Author contributions

TZ, BW, JM and GC, designed the experiments. GC, JM, ZL and BW performed laboratory work and analyzed the results. TZ, BW, JM, GC, ZL and SAK prepared the figures. GC, JM and TZ wrote the manuscript with input from all authors. All authors reviewed and approved the manuscript.

